# Regulation of neutrophil NADPH oxidase by PSTPIP2 affects bone damage in murine autoinflammatory osteomyelitis

**DOI:** 10.1101/727222

**Authors:** Jarmila Kralova, Ales Drobek, Jan Prochazka, Frantisek Spoutil, Daniela Glatzova, Simon Borna, Jana Pokorna, Tereza Skopcova, Pavla Angelisova, Martin Gregor, Pavel Kovarik, Radislav Sedlacek, Tomas Brdicka

## Abstract

Autoinflammatory diseases are characterized by dysregulation of the innate immune system leading to spontaneous inflammation. *Pstpip2*^*cmo*^ mouse strain is a well-characterized model of this class of disorders. Due to the mutation leading to the lack of adaptor protein PSTPIP2, these animals suffer from autoinflammatory chronic multifocal osteomyelitis similar to several human syndromes. Current evidence suggests that it is driven by hyperproduction of IL-1β by neutrophil granulocytes. Here we show that in addition to IL-1β, PSTPIP2 also negatively regulates ROS generation by neutrophil NADPH oxidase. *Pstpip2*^*cmo*^ neutrophils display highly elevated ROS production in response to a range of stimuli. Inactivation of NADPH oxidase in *Pstpip2*^*cmo*^ mice did not affect IL-1β levels and the autoinflammatory process was initiated with similar kinetics. However, the bone destruction was almost completely alleviated, suggesting that dysregulated NADPH oxidase activity is a key factor promoting autoinflammatory bone damage in *Pstpip2*^*cmo*^ mice.

## Introduction

Autoinflammatory diseases represent a distinct class of disorders of the innate immune system. They are characterized by a pathological inflammation that typically arises spontaneously without detectable extrinsic cause and in the absence of autoantibodies or auto-reactive T cells. The symptoms are rather diverse. The most characteristic include periodic fever attacks, skin rashes, arthralgia, myalgia, abdominal pain, arthritis, osteomyelitis, and other signs of systemic or organ specific inflammation (de Jesus, Canna, Liu, & Goldbach-Mansky, 2015; Manthiram, Zhou, Aksentijevich, & Kastner, 2017; Masters, Simon, Aksentijevich, & Kastner, 2009). A number of autoinflammatory diseases are caused by a pathological hyperactivity of IL-1β pathway, either as a result of mutations in single genes affecting inflammasomes and other components of IL-1β activation machinery, or from more complex causes, where the underlying genetic lesion is unknown (de Jesus et al., 2015; Manthiram et al., 2017).

Bone damage or other type of bone involvement are common in IL-1β-driven autoinflammatory diseases (Bader-Meunier, Van Nieuwenhove, Breton, & Wouters, 2018). IL-1β promotes osteoclast activity by stimulating RANKL expression in osteoblasts and by direct binding to osteoclasts. This way it likely stimulates inflammatory bone resorption by these cells during the course of the disease (Bader-Meunier et al., 2018; Takayanagi, 2007). Interestingly, different diseases of this group show different and distinct types of bone damage. Moreover, the bone damage is often observed only in a fraction of patients with a particular disease (Bader-Meunier et al., 2018; Houx et al., 2015; Tunca et al., 2005). These observations suggest that the character of genetic lesion, genetic modifiers or other circumstances are critically affecting the outcome (Bader-Meunier et al., 2018). However, the identity of these factors and the mechanisms of how they change the clinical picture are largely unknown.

One of the key activators of IL-1β pathway mutated in several autoinflammatory conditions is NLRP3 inflammasome. It is activated by aberrant ion fluxes, lysosomal damage by crystalline matter, such as silica or monosodium urate crystals, mitochondrial damage, presence of reactive oxygen species (ROS), as well as various PAMPs and DAMPs (Broz & Dixit, 2016; Carta et al., 2012; Gong, Yang, Jin, Jiang, & Zhou, 2018; Hughes & O’Neill, 2018; Lawlor & Vince, 2014). Several unifying mechanisms enabling recognition of such a variety of stress agents by a single type of inflammasome have been suggested, but none of them has yet gained universal acceptance (Broz & Dixit, 2016; Gong et al., 2018; Hughes & O’Neill, 2018; Lawlor & Vince, 2014; Zhen & Zhang, 2019). Production of ROS represents one such a mechanism that could connect cellular stress to NLRP3 inflammasome activation (for review see (Abais, Xia, Zhang, Boini, & Li, 2015; Carta et al., 2012)). In most of the cell types, there appear to be at least two main sources of ROS, NADPH oxidases and mitochondria (Holmström & Finkel, 2014). In phagocytes, NADPH oxidase is activated downstream of PAMPs, DAMPs and other pro-inflammatory stimuli and its products are toxic to microorganisms. Various NADPH oxidases are also part of a broad array of signaling pathways in multiple cell types (Brandes, Weissmann, & Schroder, 2014; Sumimoto, 2008). Mitochondrial ROS are produced mainly as a result of respiratory chain activity and their generation can be enhanced by stress or mitochondrial damage (Brookes, Yoon, Robotham, Anders, & Sheu, 2004; West, Shadel, & Ghosh, 2011; Zhou, Yazdi, Menu, & Tschopp, 2011).

Several studies have shown increased production of ROS in monocytes from autoinflammatory disease patients (Borghini et al., 2011; Bulua et al., 2011; Carta et al., 2015; Carta et al., 2012; Omenetti et al., 2014; Tassi et al., 2010; van der Burgh et al., 2014). In some of these works it has been proposed that these ROS are of mitochondrial origin (Bulua et al., 2011; Carta et al., 2012), but there are only limited options how to study this aspect in patients. The effects of increased ROS production, whether of mitochondrial or NADPH oxidase origin, on the development and/or severity of autoinflammatory diseases is currently unknown.

There are relatively few mouse models of autoinflammatory bone diseases. One of the best characterized is *Pstpip2*^*cmo*^ mouse strain which spontaneously develops severe bone and soft tissue inflammation mainly in hind paws and tail. In several aspects, the disease resembles a human condition known as chronic recurrent multifocal osteomyelitis (CRMO) and was thus termed chronic multifocal osteomyelitis (CMO). From there the strain derives its name *Pstpip2*^*cmo*^ (Byrd, Grossmann, Potter, & Shen-Ong, 1991). The disease is caused by a point mutation in the gene coding for the adaptor protein PSTPIP2 (Ferguson et al., 2006). As a result, no PSTPIP2 is detectable in these mice at the protein level (Chitu et al., 2009). The mechanism of how PSTPIP2 deficiency leads to CMO disease is only partially understood. It binds several inhibitory molecules, including PEST-family protein tyrosine phosphatases, phosphoinositide phosphatase SHIP1 and inhibitory kinase Csk, which likely mediate its negative regulatory effect on the inflammatory response (Drobek et al., 2015; Wu, Dowbenko, & Lasky, 1998). In addition, it has been reported that osteomyelitis in *Pstpip2*^*cmo*^ mice is completely dependent on excessive IL-1β production by neutrophilic granulocytes (S. L. Cassel et al., 2014; J. R. Lukens et al., 2014; John R. Lukens et al., 2014). Genetic studies suggest a combined involvement of the NLRP3 inflammasome and a poorly characterized mechanism dependent on caspase-8. A relatively limited role of neutrophil proteases has also been demonstrated (Gurung, Burton, & Kanneganti, 2016; John R. Lukens et al., 2014). The involvement of NLRP3 inflammasome suggests that cellular stress and ROS might be involved in CMO disease pathology, especially, when we consider the fact that neutrophils are very potent producers of NADPH-oxidase-derived ROS. Moreover, ROS are also activators of osteoclasts (Callaway & Jiang, 2015), a cell type likely responsible for inflammatory bone damage in CMO mice (Chitu et al., 2012). Here we show that in *Pstpip2*^*cmo*^ neutrophils, ROS generation is profoundly dysregulated and these cells produce substantially increased amounts of ROS in response to variety of stimuli. Strikingly, the dysregulated ROS production by these neutrophils does not have a strong effect on IL-1β production and soft tissue inflammation, but rather on the bone inflammation and subsequent bone damage, suggesting that the role of NADPH-oxidase-derived ROS is not in triggering the CMO, but rather in directing the damage accompanying this disease to the bones.

## Results

### *Pstpip2*^*cmo*^ neutrophils produce substantially more ROS in response to inflammasome activator silica than wild-type neutrophils

Disease development in *Pstpip2*^*cmo*^ mice is, in part, dependent on NLRP3 inflammasome (Gurung et al., 2016). Since ROS are involved in the NLRP3 inflammasome regulation, we tested if their production was dysregulated in *Pstpip2*^*cmo*^ bone marrow (BM) cells. We isolated these cells from wild-type C57Bl/6 (WT) and *Pstpip2*^*cmo*^ mice (backcrossed to the same genetic background) and stimulated these cells with silica particles, a well established activator of NLRP3 inflammasome (Suzanne L. Cassel et al., 2008) employed in previous studies of *Pstpip2*^*cmo*^ mice (S. L. Cassel et al., 2014; Drobek et al., 2015; J. R. Lukens et al., 2014). Strikingly, this stimulation led to a substantially stronger ROS response in *Pstpip2*^*cmo*^ cells, when compared to their WT counterparts (Figure 1A). Since ROS production is predominantly characteristic of neutrophils, which form a large fraction of bone marrow leukocytes (Figure 1B), we have isolated these cells for further testing. Silica stimulation of untouched neutrophils, isolated by negative selection, led to even higher production of ROS when compared to the full bone marrow. Moreover, the difference between WT and *Pstpip2*^*cmo*^ cells was still preserved (Figure 1C). In our experiments, negatively selected neutrophils were typically more than 90% pure. However, large fraction of the contaminating cells were monocytes. Since these cells are also known to respond by ROS production to a variety of stimuli we have analyzed ROS generation by purified monocytes. In order to determine the impact of these cells on our results we have adjusted the quantity of monocytes to 10% of the neutrophil numbers, which is similar to the amount of monocytes contaminating our neutrophil samples prepared by negative selection. We compared the response of these monocytes with the ROS production by neutrophils isolated by positive selection on Ly6G. This purification resulted in virtually pure (more than 99%) neutrophils. The response of purified monocytes was almost two orders of magnitude lower than that of purified neutrophils, and there was no significant difference between WT and *Pstpip2*^*cmo*^ cells (Figure 1D). These data demonstrated that in the neutrophil samples prepared by negative selection, the monocyte contribution to the measured ROS production is negligible. They also lead to the conclusion that even in non-separated bone marrow the vast majority of ROS originated from neutrophils, and that neutrophils are responsible for enhanced ROS production by *Pstpip2*^*cmo*^ bone marrow cells.

**Figure 1.**
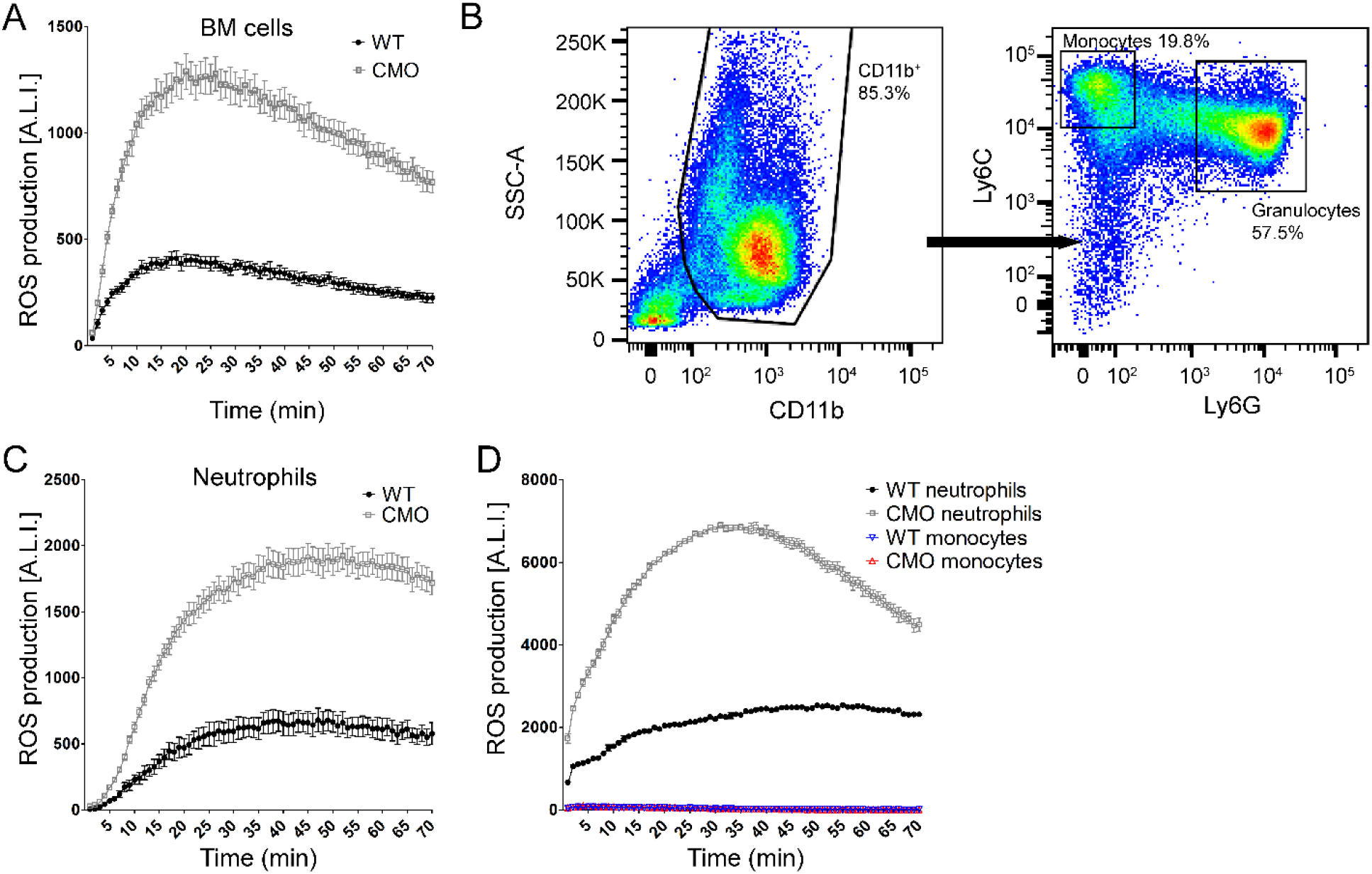
*Pstpip2*^*cmo*^ neutrophils produce higher amounts of ROS. (A) ROS production by Pstpip2^cmo^ (CMO) and wild-type (WT) bone marrow cells. BM cells (10^6^ cells per well) were treated in a 96-well plate with silica particles in the presence of 100 μM luminol. ROS-induced luminescence was measured in 1 min intervals. (B) Representative flow cytometry dot plot showing percentages of monocytes and neutrophils in Pstpip2^cmo^ BM. N > 5. (C) Similar experiment as in (A) performed on neutrophils purified by negative selection (10^6^ cells per well) (D) Similar experiment as in (A) performed on neutrophils purified by positive selection (10^6^ cells per well). The response is compared to ROS production by FACS-sorted (Ly6C^+^, Ly6G^−^) monocytes (1 × 10^5^ per well). N = 3. Representative plots are shown. In panels A, C, D, individual points and error bars represent mean ± SEM values obtained from 2-8 technical replicates. Quantification based on multiple biological replicates is shown in Figure 2 below. A.L.I. stands for Arbitrary Luminescence Intensity.

### Higher ROS production by *Pstpip2*^*cmo*^ neutrophils is observed across a range of conditions

To find out how universal the ROS overproduction in *Pstpip2*^*cmo*^ neutrophils is, we treated either BM cells or purified neutrophils with silica, PMA, live *E.coli* bacteria, heat-aggregated mouse IgG (as a model of immunocomplexes), TNF-α, LPS or fMLP. All these experiments demonstrated dysregulated ROS production in *Pstpip2*^*cmo*^ BM cells (Figure 2A-C, E) and purified neutrophils (Figure 2B, D, E). The same dysregulation was also observed in the bone marrow cells with *Pstpip2*^*cmo*^ mutation on Balb/c genetic background (Figure S1). These results show that PSTPIP2 deficiency renders neutrophils more sensitive and prone to produce more ROS than WT cells. Interestingly, unstimulated BM cells from *Pstpip2*^*cmo*^ mice produced low but detectable levels of ROS even in the absence of any stimulus. This constitutive production has not been observed in WT BM (Figure 2A, F).

**Figure 2.**
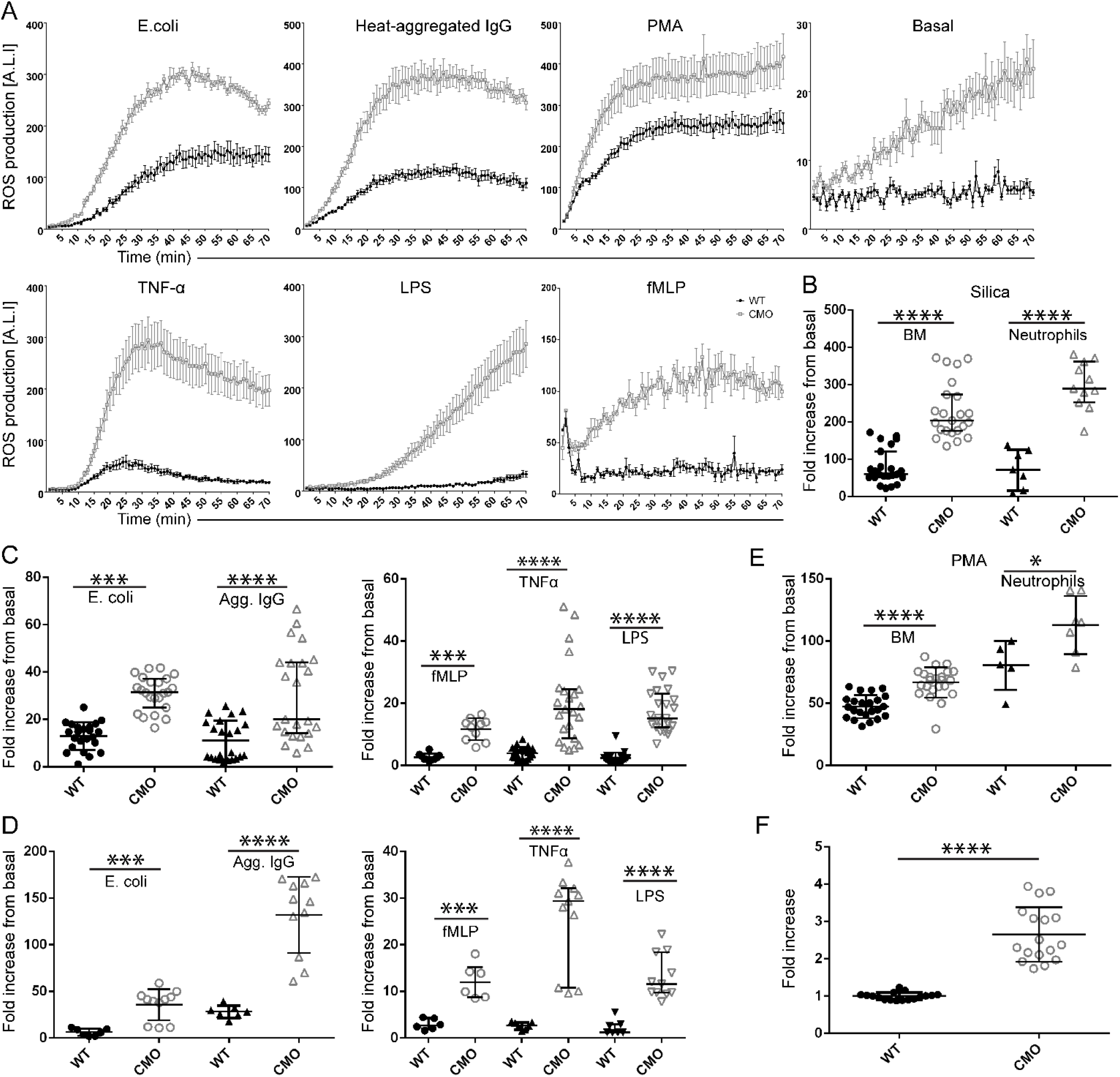
Increased ROS production by *Pstpip2*^*cmo*^ neutrophils is triggered by a variety of stimuli. (A) ROS production by Pstpip2^cmo^ (CMO) and wild-type (WT) bone marrow cells treated with indicated agents (E.coli culture OD600 = 0.8, diluted 1:5; 300 μg/ml heat-agregated murine IgG (Agg. IgG); 100 ng/ml PMA; 10 ng/ml TNFα; 100 ng/ml LPS; 1 μg/ml fMLP). Graphs from representative experiments are shown with the data depicted as mean ± SEM from 2-3 technical replicates. (B) Quantification of ROS production by BM cells and neutrophils after silica treatment. (C) Quantification of ROS production by BM cells treated with live E.coli bacteria (E.coli), mouse aggregated IgG (IC), fMLP, TNFα, and LPS. (D) Similar analysis as in (C), performed on neutrophils purified by negative selection. (E) Quantification of ROS production by BM cells and neutrophils after treatment with PMA. (F) Quantification of basal ROS production by non-stimulated BM cells. Fold increase (B-E) was calculated as a ratio between maximum response after stimulation and basal ROS production of unstimulated cells in each experiment. In (F), fold increase is calculated as a ratio of maximum response and mean ROS production by WT cells. In (B-F), data points represent results from biological replicates, each of them mean of 2-3 technical replicates. Bars represent median with interquartile range. See also Figure S1.

### Excessive ROS production by *Pstpip2*^*cmo*^ neutrophils is not a consequence of ongoing inflammation

Higher ROS production under resting conditions and after activation with a wide range of stimuli demonstrates general dysregulation of pathways leading to ROS production in *Pstpip2*^*cmo*^ neutrophils. This dysregulation could be cell intrinsic due to PSTPIP2 deficiency or a side effect of ongoing bone inflammation, which could prime bone marrow neutrophils located in the proximity of the inflamed tissue. It has previously been reported that autoinflammation in *Pstpip2*^*cmo*^ mice is completely dependent on IL-1β and its receptor (S. L. Cassel et al., 2014; J. R. Lukens et al., 2014). Signaling through this receptor is critically dependent on MyD88 adaptor protein (Adachi et al., 1998). In order to determine whether the observed overproduction of ROS in *Pstpip2*^*cmo*^ neutrophils is not just the effect of ongoing inflammation, we crossed *Pstpip2*^*cmo*^ mice with MyD88-deficient strain to block IL-1β signaling. As expected, *Pstpip2*^*cmo*^ mice were in the absence of MyD88 completely protected from the disease development as determined by visual inspection (Figure 3A) and X-ray micro computed tomography (X-ray μCT) of hind paws (Figure 3B). MyD88-deficient *Pstpip2*^*cmo*^ bone marrow cells displayed the same dysregulation in ROS production triggered by a variety of stimuli as *Pstpip2*^*cmo*^ cells. Treatments with LPS or *E.coli* were the only exceptions where the response was lower in both *Pstpip2*^*cmo*^/*MyD88*^−/−^ and *MyD88*^−/−^ cells, probably due to the higher dependence of signaling triggered by these activators on the TLR/MyD88 pathway. On the other hand, even in *MyD88*-deficient cells the *cmo* mutation gave rise to a stronger response to these two stimuli when compared to *MyD88* deficient cells without the *cmo* mutation (Figure 3C). These results demonstrate the cell intrinsic dysregulation of NADPH oxidase machinery that is not caused by chronic exposure to the inflammatory environment.

**Figure 3.**
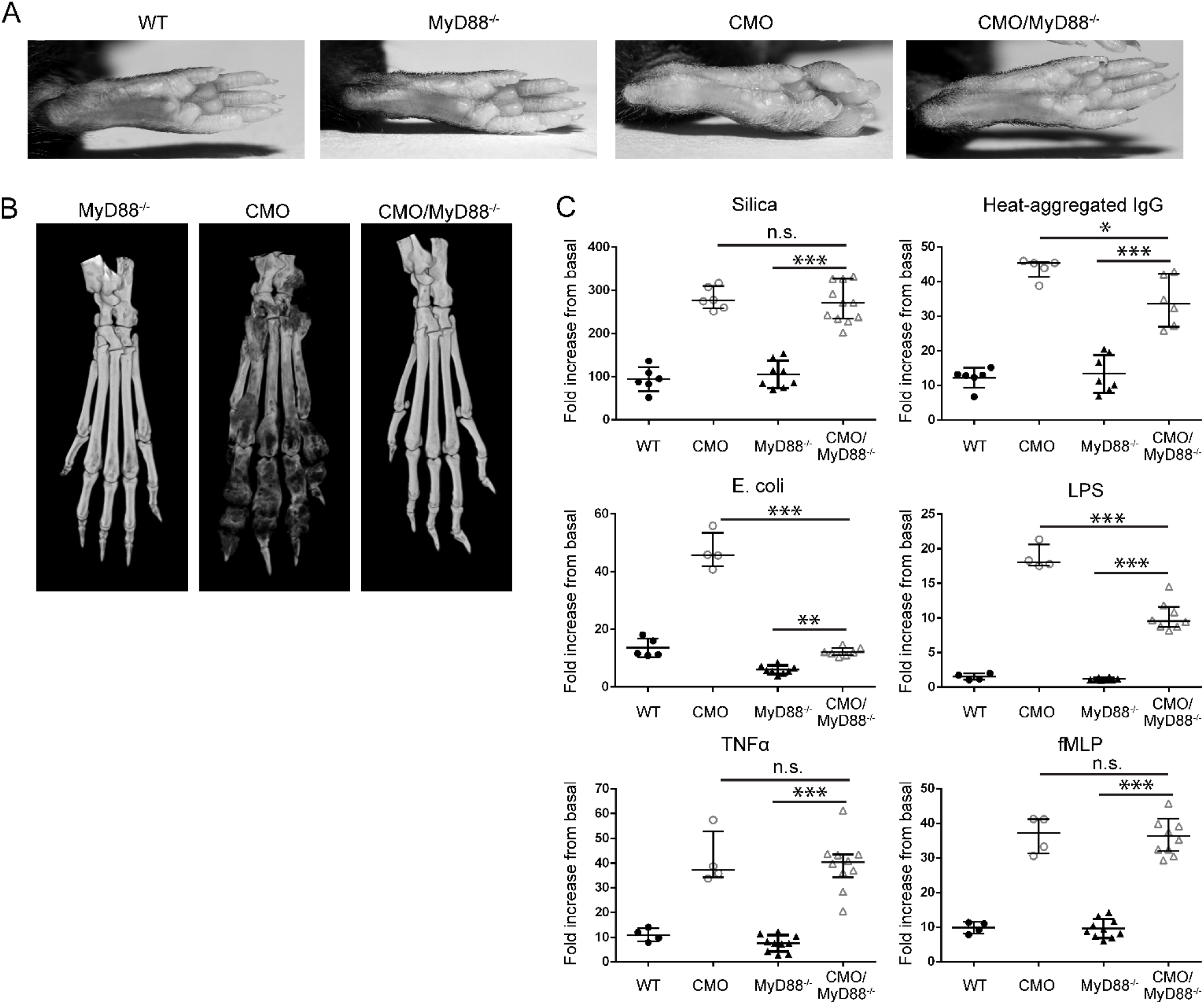
Overproduction of ROS is not triggered by ongoing inflammation. (A) Representative hind paw photographs of 23 - 26 week old WT, MyD88^−/−^, Pstpip2^cmo^ (CMO) and Pstpip2^cmo^/MyD88^−/−^ (CMO/MyD88^−/−^) mice. For Pstpip2^cmo^/MyD88^−/−^, 24 animals between 13 and 34 weeks of age were photographed, none of which showed any symptom of CMO disease. (B) Representative X-ray μCT scans of hind paws from MyD88^−/−^, Pstpip2^cmo^ and Pstpip2^cmo^/MyD88^−/−^ mice, 14 weeks old. N = 5 for Pstpip2^cmo^/MyD88^−/−^ and N = 3 for MyD88^−/−^ (C) Quantification of ROS production by BM cells isolated from WT, MyD88^−/−^, Pstpip2^cmo^ and Pstpip2^cmo^/MyD88^−/−^ mice and treated with indicated stimuli. Fold increase was calculated as a ratio between maximum response after stimulation and basal ROS production of unstimulated cells in each experiment. Data points represent results from biological replicates (individual mice), each of them mean of 2-3 technical replicates. Bars represent median with interquartile range.

### ROS hyperproduction in *Pstpip2*^*cmo*^ neutrophils is suppressed by PSTPIP2 binding partners and is accompanied by hyperphosphorylation of p47-phox

To elucidate the mechanism of how PSTPIP2 suppresses ROS production, we employed conditionally immortalized *Pstpip2*^*cmo*^ granulocyte progenitors (Wang et al., 2006) we had established previously (Drobek et al., 2015) and reconstituted these cells with doxycycline-inducible retroviral constructs coding for WT PSTPIP2 and its mutated versions unable to bind PEST-family phosphatases (W232A) and SHIP1 (3YF) (Drobek et al., 2015; Wu et al., 1998). After maturation of these progenitors into neutrophils and induction of PSTPIP2 expression with doxycycline, we treated these cells with silica and measured ROS response. We observed around 50% reduction of ROS response in cells expressing WT PSTPIP2. In contrast, both mutated versions of PSTPIP2 were unable to substantially inhibit silica-induced ROS generation, despite similar expression levels of these constructs (Figures 4A and S2).

**Figure 4.**
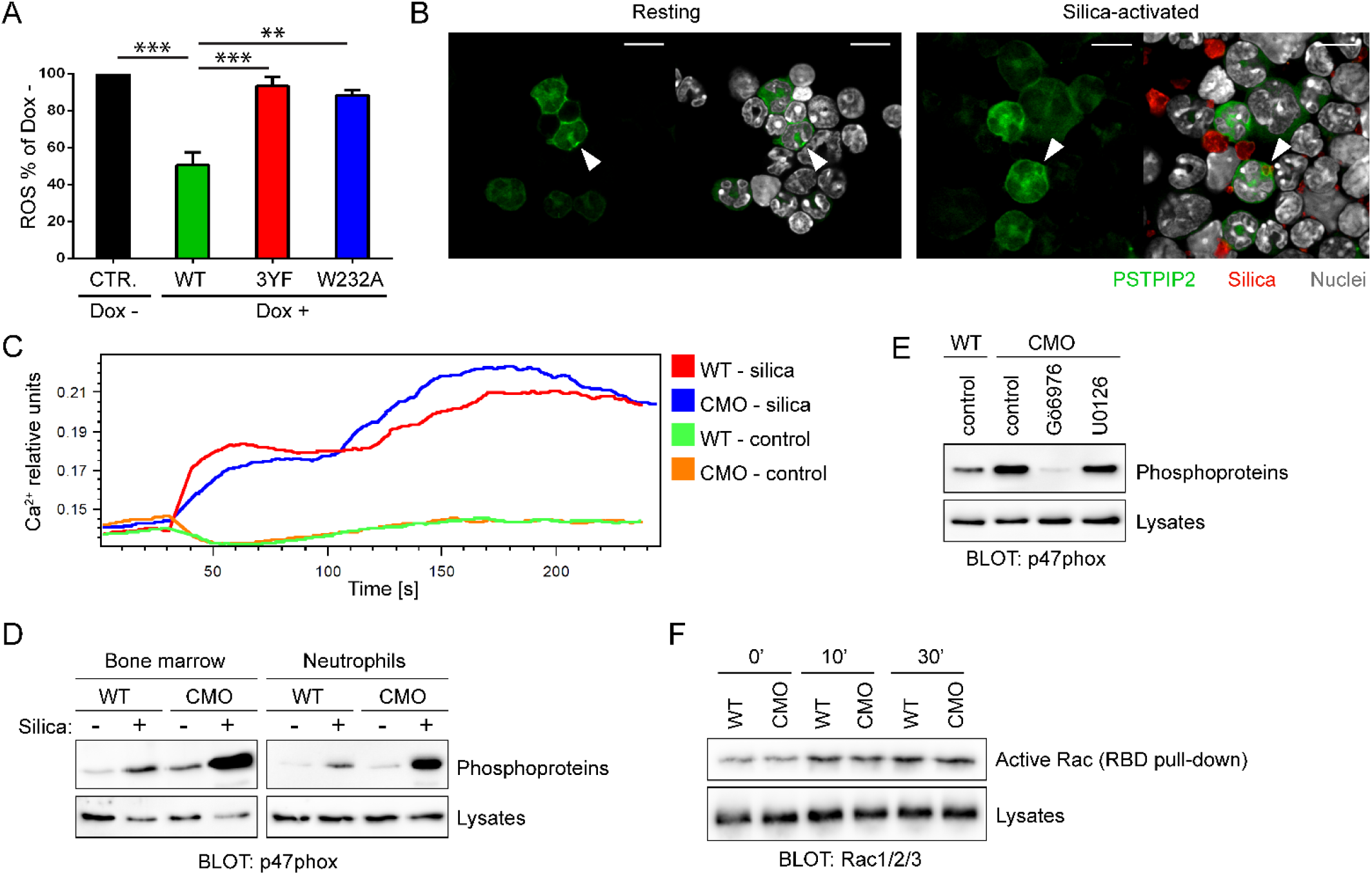
PSTPIP2-mediated suppression of ROS production in neutrophils is dependent on PSTPIP2 binding partners and involves negative regulation of p47phox phosphorylation. (A) Suppression of ROS production by wild-type and mutant PSTPIP2 in mature granulocytes differentiated from conditionally immortalized granulocyte progenitors transduced with inducible PSTPIP2 constructs. Expression of PSTPIP2 was triggered by overnight incubation with doxycycline. On the vertical axis percentage of ROS response by Dox treated cells when compared to Dox non-treated cells is shown. Error bars represent SEM of 3 biological replicates. Only significant differences are highlighted with asterisks. (B) Subcellular localization of PSTPIP2 in resting and silica-activated neutrophils differentiated in vivo from bone marrow cells transduced with retroviral PSTPIP2-EGFP construct. Bar = 10 μm. N = 4. (C) Calcium response of WT and Pstpip2^cmo^ neutrophils (gated from full BM) after silica treatment. N = 3 (D) Phosphorylation of p47phox in WT and Pstpip2^cmo^ neutrophils after 30 min incubation with silica. To detect p47phox phosphorylation, total phosphoproteins were isolated from neutrophil lysates followed by p47phox detection by immunoblotting. N = 3 (E) Phosphorylation of p47phox in Pstpip2^cmo^ neutrophils treated for 30 min with silica together with PKC inhibitor Gö6976 or MEK inhibitor U0126. Phosphorylation of p47phox was detected as in (D). N = 4 for Gö6976 and N = 2 for U0126. (F) RAC activation in WT and Pstpip2^cmo^ neutrophils. Active RAC was isolated using PAK1-RBD-GST and Glutathion-Sepharose and detected by immunoblotting with antibody to all three RAC proteins (RAC1/2/3). N = 2. See also Figure S2.

To analyze subcellular localization of PSTPIP2 during silica stimulation we isolated bone marrow progenitors from *Pstpip2*^*cmo*^ mice and transduced these cells with retroviral construct coding for PSTPIP2 fused to EGFP. Next, we transplanted these cells into lethally irradiated mice and after 2 weeks we collected neutrophils expressing PSTPIP2-EGFP for microscopy analysis. In neutrophils, PSTPIP2 showed diffuse distribution throughout the cytoplasm with occasional formation of speckles in a small fraction of cells (Figure 4B, left panel, see an arrowhead). After addition of fluorescently labelled silica particles, neutrophils interacted with these particles and phagocytosed some of them (Figure 4B, right panel, see an arrowhead). However, we did not observe any changes in PSTPIP2 subcellular localization during this process (Figure 4B). This result suggests that large-scale redistribution of PSTPIP2 inside the cells is not part of the mechanism of how PSTPIP2 controls neutrophil activity during the treatment with silica.

In order to identify the dysregulated process leading to ROS overproduction at the biochemical level, we measured the calcium response in WT and *Pstpip2*^*cmo*^ BM cells. Cells were loaded with Fura Red dye and stimulated with silica particles. We observed the same calcium response in both WT and *Pstpip2*^*cmo*^ cells (Figure 4C) indicating that proximal signaling steps leading to calcium response are not responsible for increased ROS production in *Pstpip2*^*cmo*^ cells.

One of the major events further downstream is phosphorylation of NADPH oxidase cytosolic subunits by members of protein kinase C (PKC) family, including phosphorylation of p47phox, which then serves as an assembly hub for building the active NADPH oxidase complex (Brandes et al., 2014). To detect p47phox phosphorylation we isolated phosphoproteins from untreated and silica treated BM cells or purified neutrophils and detected p47phox in the isolated material by immunoblotting. In both *Pstpip2*^*cmo*^ BM cells and neutrophils we found substantially stronger phosphorylation of p47phox when compared to WT cells (Figure 4D).

In addition to the PKCs, p47-phox can also be phosphorylated by other kinases, such as Erk, which is dysregulated in silica-treated CMO neutrophils (Dang et al., 2006; Drobek et al., 2015). To identify the kinases involved in the observed hyperphosphorylation of p47phox, we treated *Pstpip2*^*cmo*^ neutrophils either with PKC inhibitor Gö6976 or MEK1/MEK2 inhibitor U0126 prior to activation with silica. Only the treatment with PKC inhibitor led to specific block of p47-phox phospholylation (Figure 4E).

Small G-protein RAC is another critical component of active NADPH oxidase. We have tested the activation status of RAC after silica treatment of bone marrow cells, but no difference between WT and *Pstpip2*^*cmo*^ cells has been observed (Figure 4F)

Collectively these data suggest that PSTPIP2 via its binding partners, suppresses pathways leading to PKC-mediated p47-phox phosphorylation and that this is the mechanism how PSTPIP2 attenuates NADPH oxidase activity and ROS production.

### Unprovoked ROS production by neutrophils *in vivo* precedes the onset of the disease

To analyze the ROS production in vivo during the disease development we used luminol derivative L-012 to visualize ROS generation in living anesthetized mice. Very interestingly, we observed a strong luminescent signal already in freshly weaned 3 week old mice that were otherwise asymptomatic (Figure S3). The signal was mostly localized along the tail and with weaker intensity in the hind paws. Visualization at later time points revealed that at 4 weeks of age the ROS production was equally intensive in the tail and paws (Figure 5A, B) and gradually moved to the hind paws during the weeks 6-8. At this age, ROS production became predominant in hind paws with more restricted focal localization (Figures 5B and S3).

**Figure 5.**
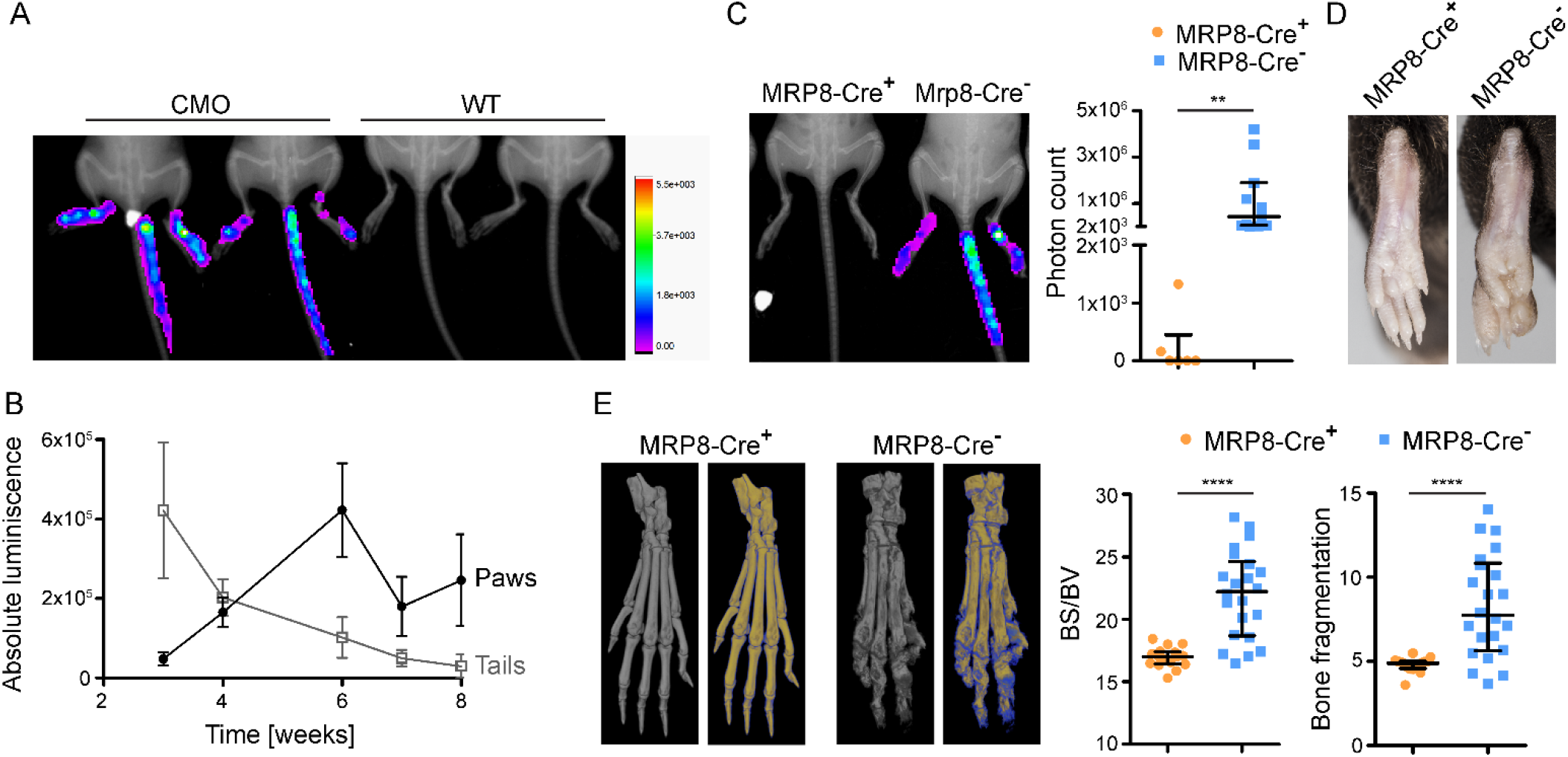
Neutrophil-dependent ROS production in vivo precedes the onset of the disease. (A) Representative in vivo measurement of ROS in 4 week old anesthetized WT and Pstpip2^cmo^ (CMO) mice. Mice were injected intraperitoneally with L-012 (75 mg/kg) and ROS-induced luminescence was measured in the whole body imager. The luminescence is shown as a heat-map in artificial colors on the background of an X-ray image. (B) Time course quantification of absolute L-012 luminescence driven by in vivo ROS production in Pstpip2^cmo^ mice. The luminescence was quantified separately for paws and tails. N = 6 per time point, mean ± SEM. (C) Representative in vivo measurement of ROS production in anesthetized 4 week old Pstpip2^cmo^-DTA mice where neutrophils were depleted via MRP8-CRE-dependent expression of Diphtheria toxin (MRP8-CRE^+^) or left untouched in the absence of MRP8-CRE (MRP8-CRE^−^). The left panel shows quantification of in vivo ROS production in multiple mice of both genotypes (3-4 weeks old, sex and age matched). (D) Representative photographs of hind paws of 26 week old mice of the same genotypes as in (C) (E) Representative X-ray μCT scans of bones from 20 week old mice of the same genotypes as in (C). Grey images represent visualization of total bone tissue. Pseudocolor images distinguish between old (in yellow) and newly formed (in blue) bone mass. Quantification of bone surface/bone volume ratio (BS/BV) and bone fragmentation in paw bones of multiple mice is shown on the left. Bars in (C,E) represent median with interquartile range. See also Figure S3.

To test if neutrophils are the source of dysregulated ROS observed *in vivo*, we have generated *Pstpip2*^*cmo*^ mouse strain where majority of neutrophils were deleted via MRP8-CRE-dependent expression of Diphtheria toxin (*Pstpip2*^*cmo*^*-DTA-MRP8-CRE*, Figure S3). *In vivo* ROS imaging revealed that ROS production in the tails and hind paws of these mice was almost completely abolished (Figure 5C). In addition, these mice also did not show any symptoms of autoinflammatory disease, whether determined by visual inspection (Figure 5D) or by X-ray μCT analysis (Figure 5E). These data strongly support the idea that increased ROS production preceding the onset of the disease originates in neutrophils and, at the same time, confirm that neutrophils are critical for the development of disease symptoms (John R. Lukens et al., 2014).

### ROS deficiency has specific effects on bone destruction

Strong unprovoked production of ROS in very young mice preceding visible symptoms weeks before their demonstration suggested that ROS may act upstream of IL-1β in osteomyelitis development. To determine the contribution of high *in vivo* ROS generation to disease development, we crossed *Pstpip2*^*cmo*^ mice to gp91phox-deficient mouse strain. In the absence of gp91phox, we were unable to detect any ROS production even in *Pstpip2*^*cmo*^ cells (Figures 6A and S4). These data confirm that NADPH oxidase was responsible for the dysregulated ROS production in *Pstpip2*^*cmo*^ neutrophils. Surprisingly, *Pstpip2*^*cmo*^ mice lacking gp91phox developed similar disease symptoms like *Pstpip2*^*cmo*^ mice (Figure 6B) and with similar, only slightly delayed, kinetics (Figure 6C). Blind scoring of the disease severity by visual inspection of hind paw photographs collected throughout various experiments revealed that the symptoms of the disease are only partially alleviated in gp91phox-deficient animals, approximately by 1-2 points on 8-point scale (Figure 6D). Moreover, ELISA analysis detected comparable amount of IL-1β in hind paw extracts from *Pstpip2*^*cmo*^ and *Pstpip2*^*cmo*^/*gp91phox*^−/−^ animals (Figure 6E) and similar amount of processed IL-1β p17 was found in the lysates of silica-stimulated bone marrow cells by immunoblot (Figure 6F).

**Figure 6.**
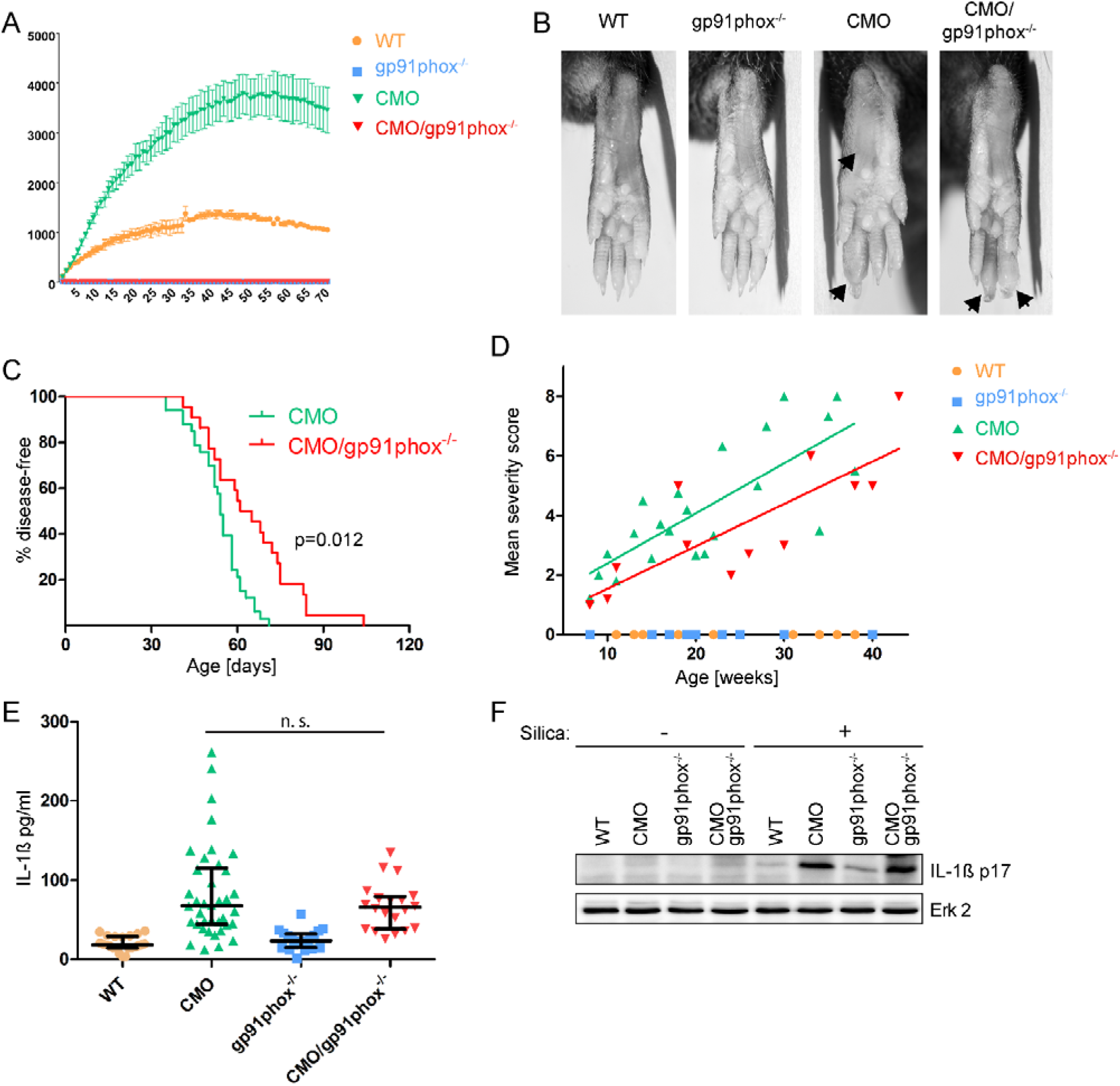
Limited effect of gp91phox deficiency on visible symptom development and IL-1β production. (A) In vitro ROS response induced by silica treatment in BM cells from WT, Pstpip2^cmo^, gp91phox^−/−^ and Pstpip2^cmo^/gp91phox^−/−^ mice. (B) Representative photographs of hind paws of 18-19 week old WT, Pstpip2^cmo^, gp91phox^−/−^ and Pstpip2^cmo^/gp91phox^−/−^ mice. (C) Disease-free curve comparing the time of disease appearance in Pstpip2^cmo^ and Pstpip2^cmo^/gp91phox^−/−^ mice. Development of visible symptoms was evaluated 2-3 times per week. (D) Disease severity was scored (scale from 0 to 8) by visual inspection of photographs of the hind paws collected over the course of this study. Each point represents mean value for the mice of the same age and genotype. Lines were generated using linear regression. (E) ELISA analysis of IL-1β from hind paw lysates. Samples were adjusted to the same protein concentration before analysis. Bars represent median with interquartile range. (F) Presence of processed IL-1β in the lysates from bone marrow cells treated for 60 min (or not) with silica was detected by immunoblotting. N = 4. See also Figure S4.

These data demonstrated that the phagocyte NADPH-oxidase is dispensable for autoinflammatory disease initiation, but it affects the severity of the disease. We also noticed that the character of the hind paw edema was somewhat different in *Pstpip2*^*cmo*^/*gp91phox*^−/−^ mice. Typically, the swelling was most serious at the distal part of phalanges and only rarely affected metatarsal area in *Pstpip2*^*cmo*^/*gp91phox*^−/−^ animals, while in *Pstpip2*^*cmo*^ mice metatarsi were frequently enlarged and the phalanges were often most seriously affected in their central parts (Figure 6B).

In order to find out if these differences were caused by different character of bone inflammation we performed X-ray μCT analysis of *Pstpip2*^*cmo*^ and *Pstpip2*^*cmo*^/*gp91phox*^−/−^ mice. Very surprisingly, bone destruction in *Pstpip2*^*cmo*^/*gp91phox*^−/−^ animals was almost entirely missing, while in *Pstpip2*^*cmo*^ mice substantial bone damage could be observed (Figure 7A). To support this observation with a quantitative analysis, we calculated bone surface to volume ratio and bone fragmentation from the X-ray μCT data. *Pstpip2*^*cmo*^/*gp91phox*^−/−^ mice showed similar values to WT, while values for *Pstpip2*^*cmo*^ mice were substantially higher (Figure 7B). Timeline X-ray μCT scans of hind paws revealed progressive bone lesion formation in *Pstpip2*^*cmo*^ mice while *Pstpip2*^*cmo*^ *gp91phox*^−/−^ littermates remained largely protected (Figure S5).

**Figure 7.**
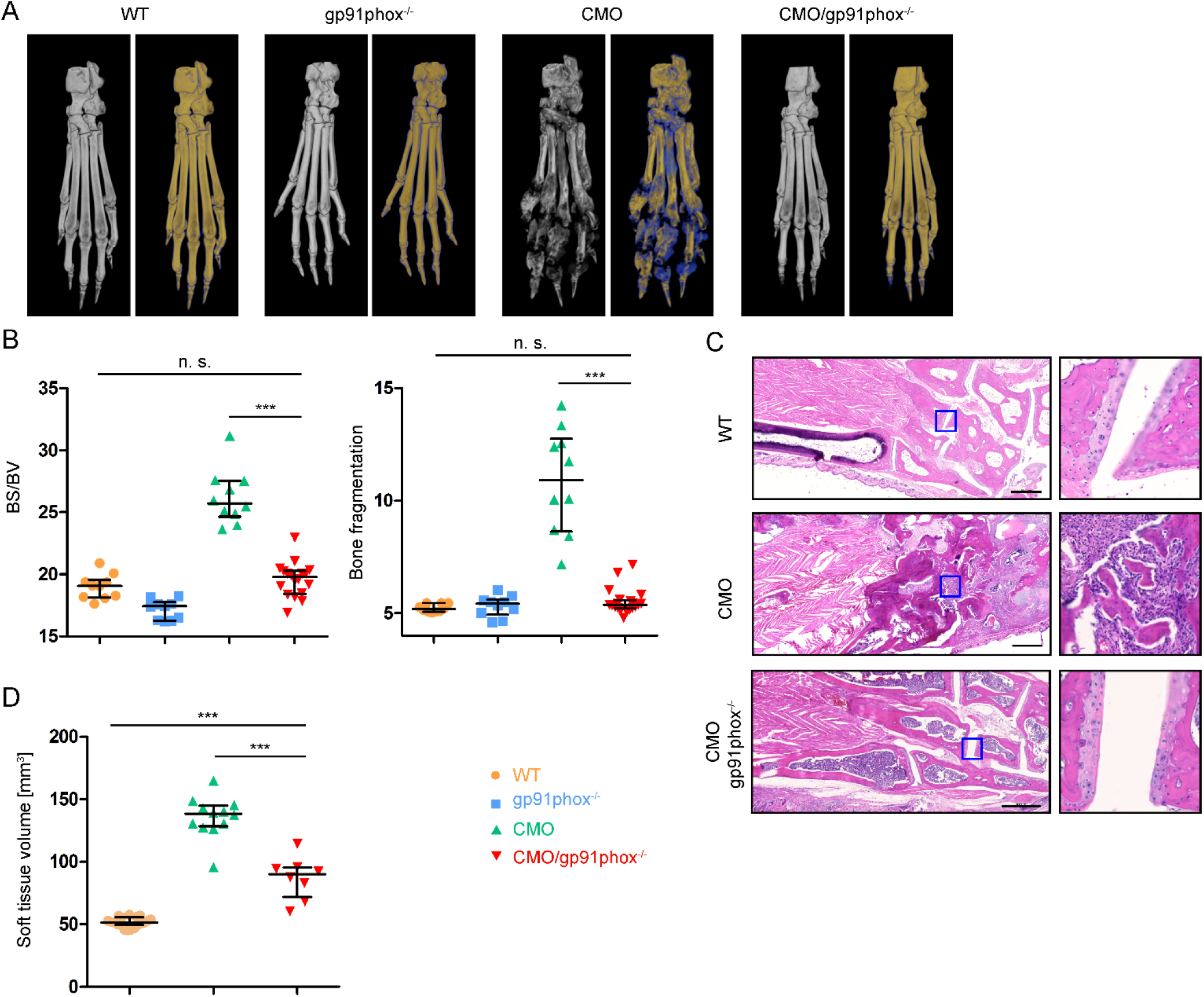
Almost complete absence of bone damage in gp91phox-deficient *PSTPIP2*^*cmo*^ mice. (A) Representative X-ray μCT scans of bones from 14 week old mice. Grey images represent visualization of total bone tissue. Pseudocolor images distinguish between old (in yellow) and newly formed (in blue) bone mass. (B) Quantification of bone surface/bone volume ratio (BS/BV) and bone fragmentation in paw bones of 14 weeks old mice. Bars represent median with interquartile range. (C) Sections of paraffin embedded tissue from tarsal area of hind paw stained with hematoxylin-eosin. (D) Volume of soft tissue in hind paws calculated as a total paw volume from which the bone volume has been subtracted. Values were calculated from X-ray μCT data. Bars represent median with interquartile range. See also Figure S5.

The lack of bone damage can also be demonstrated on tissue sections from tarsal area of hind paws (Figure 7C). The CMO mice show very high level of osteolysis of tarsal bones with almost missing joint cartilages due to arthritic changes accompanied with robust granulomatous infiltration. The WT mice have normally developed and structurally well-defined tarsal bones with undamaged joint cartilages with no infiltration of immune cells and no adverse changes in the bone marrow. The *Pstpip2*^*cmo*^/*gp91phox*^−/−^ mice showed rescue effect in ossified parts of tarsal bones with no or minimal signs of bone damage by immune cells, the soft tissue infiltration is in metatarsal area minimal compared to CMO mice. The cartilages are also well shaped and are covering the joint areas comparably with WT animals. The difference from WT animals is in hypercellular structure of bone marrow resulting in decreased volume of ossified tissue.

To address protection potential of gp91phox deficiency in old *Pstpip2*^*cmo*^ mice we performed X-ray μCT scans on 7 month old mice. Old *Pstpip2*^*cmo*^ mice suffered from strong bone destruction and remodeling but *Pstpip2*^*cmo*^/*gp91phox*^−/−^ mice were still protected from adverse effects of osteomyelitis (Figure S5). To gain a quantitative insight into the level of soft tissue inflammation we have performed computational reconstruction of soft tissues from X-ray μCT scans described above in Figure 7B and calculated soft tissue volume. This measurement revealed that soft tissues in *Pstpip2*^*cmo*^/*gp91phox*^−/−^ hind paws were significantly enlarged, albeit not to the same extent as in *Pstpip2*^*cmo*^ mice (Figure 7D). Collectively, these data demonstrate that despite significant swelling that can be detected in the hind paw soft tissues of *Pstpip2*^*cmo*^/*gp91phox*^−/−^ mice, bones remain largely protected in the absence of NADPH oxidase activity.

## Discussion

Monogenic autoinflammatory diseases develop as a result of dysregulation of the innate immune system. Although the specificity of this branch of the immune system is relatively limited, these diseases still show tissue and organ selectivity. The mechanisms of this selectivity are often poorly understood (de Jesus et al., 2015). *Pstpip2*^*cmo*^ mice represent an important model of tissue-selective IL-1β-driven autoinflammatory disease that affects mainly bones and surrounding tissue in hind paws and tails (Byrd et al., 1991; Chitu et al., 2009; Ferguson et al., 2006). Our current studies demonstrate that IL-1β pathway is not the only pathway dysregulated in these animals. ROS production by neutrophil NADPH oxidase is also substantially enhanced, independently of IL-1β activity. Moreover, our data suggest that dysregulated ROS production is a critical part of the selectivity mechanisms governing specific damage to the bones.

We found that increased ROS production in *Pstpip2*^*cmo*^ mice was associated with p47phox hyperphosphorylation, leading to the conclusion that PSTPIP2 is a negative regulator of p47phox. This phosphorylation could be almost completely abolished by inhibition of Protein kinases C (PKC). These kinases are known to be activated by calcium ions and diacylglycerol, which are both produced as a result of phospholipase C (PLC) activity (Kadamur & Ross, 2013; Lipp & Reither, 2011). The calcium response, which is a hallmark of PLC activity appeared normal in silica-stimulated *Pstpip2*^*cmo*^ neutrophils, where the scale of ROS dysregulation was largest. Therefore, we assume that the signaling is altered either at the level of PKC regulation that is independent of PLC activity, or that PSTPIP2 controls an unknown negative regulator of p47phox phosphorylation (e.g. a phosphatase) that is independent of PKCs. Further experiments are needed to clarify this issue and determine individual components of the pathway connecting PSTPIP2 to p47phox phosphorylation.

To our knowledge, genetically-determined hyperactivation of NADPH oxidase has not been described or studied in the context of IL-1β activation or in autoinflammatory disease yet. Our observations show that elevated NADPH oxidase ROS do not affect IL-1β pathway but rather the inflammatory bone damage in CMO. These data are in agreement with several other reports that disprove the role of NADPH oxidase-generated ROS in IL-1β processing by inflammasome. They are mainly based on analyses of monocytes and macrophages from NADPH-oxidase-deficient patients and mice, where IL-1β production is not altered or it is even enhanced (Meissner, Molawi, & Zychlinsky, 2008; Meissner et al., 2010; van Bruggen et al., 2010; van de Veerdonk et al., 2010). In a single study on human neutrophils, NADPH oxidase deficiency also did not lead to reduction of NLRP3 inflammasome activity (Gabelloni et al., 2013). On the other hand, inhibition of mitochondrial ROS production in monocytes/macrophages results in an impairment of IL-1β production in these cells (Nakahira et al., 2011; Zhou et al., 2011), showing that the majority of ROS supporting inflammasome activation in these cell types is generated by mitochondria.

The roles of IL-1β and ROS in CMO pathophysiology appear to be different form each other. While dysregulated IL-1β production is a critical trigger of the disease development, enhanced ROS production modifies the outcome. However, ROS are not able to initiate the disease on their own in the absence of IL-1β signaling. It is documented by our experiments with *MyD88*-deficient *Pstpip2*^*cmo*^ mice, which displayed the same ROS dysregulation as *Pstpip2*^*cmo*^ mice and yet they did not develop any symptoms of autoinflammation. These data also demonstrate that enhanced ROS production is not downstream of IL-1β, since MyD88 is critical for signaling by IL-1 receptor (Deguine & Barton, 2014).

ROS are known to play a key role in differentiation and activity of osteoclasts. These cells are responsible for physiological bone resorption during bone remodeling processes. They are also involved in pathological bone damage in a number of disease states (Okamoto et al., 2017). *Pstpip2*^*cmo*^ mice exhibited increased osteoclastogenesis and osteoclast hyperactivity suggesting that osteoclasts are responsible for inflammatory bone damage in these mice (Chitu et al., 2012). Our data show that the bone damage can be almost completely abolished when phagocyte NADPH oxidase is inactivated by deletion of its gp91phox subunit. One possibility is that deficiency in osteoclast gp91phox results in defects in their differentiation and activity and reduced bone damage. However, in *gp91phox*^−/−^ mice no bone abnormalities have been observed. In addition, gp91phox-deficient osteoclasts differentiate normally and have normal bone resorption activity (Lee et al., 2005; Sasaki et al., 2009; Yang, Madyastha, Bingel, Ries, & Key, 2001). These results show that gp91phox expressed in osteoclasts is dispensable for differentiation and activity of these cells. In fact, other NADPH oxidases were shown to be more important for their function (Goettsch et al., 2013; Lee et al., 2005). On the other hand exogenous ROS generated in culture media after addition of xanthine oxidase were shown to upregulate osteoclast numbers and activity in bone cultures in vitro (Garrett et al., 1990). Our data together with published results thus favor the explanation that exogenous ROS originating from hyperactive neutrophils, ample production of which we observed in *Pstpip2*^*cmo*^ mice *in vivo*, lead either directly or indirectly to increased differentiation and/or activity of osteoclasts and resulting bone damage.

PSTPIP2 mutations in humans have not yet been described. However, PSTPIP2 gene has been sequenced only in a limited number of CRMO patients and patients with closely related SAPHO syndrome (Ferguson et al., 2008; Hurtado-Nedelec et al., 2010; Jansson et al., 2007). CRMO and SAPHO form a rather heterogeneous disease spectrum, which may in fact represent a number of distinct disorders where various defects at the molecular level may lead to similar outcome, and so PSTPIP2 mutations in some of these patients may still be discovered in the future. The data on ROS production in these diseases are also largely missing. We are aware of only a single study where the ROS production by neutrophils was analyzed in two SAPHO patients from a single family without any mutations in PSTPIP2 gene. These data showed reduced ROS response after activation with multiple activators, including PMA, fMLP and TNFα when compared to healthy controls (Ferguson et al., 2008). However, from the information provided it was unclear whether the patients were undergoing anti-inflammatory treatment that could suppress the response at the time of analysis. Further studies are needed to fully understand the role of PSTPIP2 and ROS in CRMO, SAPHO and other inflammatory bone diseases in humans.

Inflammatory bone damage is a serious problem accompanying a number of human disorders. Full understanding of possible mechanisms that can govern its development is critical for designing successful therapeutic interventions. Our data reveal how dysregulated ROS production results in bone damage in the specific case of CMO. However, these findings may represent a more general mechanism with broader validity for other syndromes where inflammatory bone damage is involved and analysis of ROS production in other instances of inflammatory bone damage may prove beneficial.

## Materials and methods Antibodies

Antibodies to the following murine antigens were used: RAC1/2/3 (Cell Signaling Technology, Danvers, MA); p47, Erk2 (Santa Cruz Biotechnology, Dallas, TX); B220-biotin, TER119-biotin, c-Kit-biotin, CD3ɛ-biotin, Ly6G-biotin, CD115-biotin, CD11b-APC, CD11b-FITC, B220-FITC, Ly6C-PE-Cy7, Ly6C-FITC, Ly6G-FITC, Ly6G-APC, c-Kit-PE, Sca-1-APC, CD16/32-PE/Cy7 (<u>Biolegend</u>, San Diego, CA); CD34-FITC, DX5-biotin, F4/80-biotin, Thy1.2-FITC (<u>eBioscience</u>, ThermoFisher, Waltham, MA); Fc Bloc (2.4G2) (BD Biosciences, San Jose, CA); HRP-conjugated goat anti-mouse IgG (Sigma), HRP-goat anti-rabbit (Bio-Rad, Hercules, CA). The mouse mAb that recognizes murine PSTPIP2 has been described earlier (Drobek et al., 2015). Heat aggregated IgG was prepared as follows: IgG was purified from mouse serum (Sigma-Aldrich) on protein A-Sepharose (GE Healthcare, Uppsala, Sweden), transferred to PBS, and concentrated to 30 mg/ml on an Amicon Ultracel–30K unit (Millipore, Merck, Darmstadt, Germany). The aggregation was induced by heating to 63°C for 30 min.

### Other reagents

In this study we also used Luminol, Lipopolysaccharides from Escherichia coli O127:B8, fMLP, PMA (all from Sigma-Aldrich), L-012 (Wako Chemicals, USA), TNFα, G-CSF (Peprotech, Rocky Hill, NJ), U0126 (Cell Signalling Technology), Gö6976 (Calbiochem, Merck). Silica (silicon dioxide crystals) was obtained from Sigma. To enable fluorescent labelling for microscopy, 5 mg/ml silica particles were first coated with non-fat dry milk (2% in PBS, 1 h at room temperature) and then labelled with 5 μM Cell Proliferation Dye eFluor 670 (eBioscience), 30 min at 37°C.

### Mice

*Pstpip2*^*cmo*^ mouse strain (C.Cg-*Pstpip2*^*cmo*^/J) carrying the c.293T→C mutation in the Pstpip2 gene (on Balb/C genetic background) resulting in an L98P change in the PSTPIP2 protein (Byrd et al., 1991; Ferguson et al., 2006), B6.129S-*Cybb*^*tm1Din*^/J lacking NADPH oxidase subunit gp91phox (Pollock et al., 1995), Myd88-deficient mouse strain (B6.129P2(SJL)-*Myd88*^*tm1.1Defr*^/J, derived from *MyD88*^fl^ mice (Hou, Reizis, & DeFranco, 2008), B6.SJL-Ptprca Pepcb/BoyJ (CD45.1^+^) congenic strain (Shen et al., 1985), *B6.Cg-Tg(S100A8-cre,-EGFP)1Ilw*/*J* with granulocyte-specific CRE expression (MRP8-Cre) (Passegue, Wagner, & Weissman, 2004), and *Gt(ROSA)26Sor*^*tm1(DTA)Lky*^/*J* strain (Voehringer, Liang, & Locksley, 2008), where Diphtheria toxin expression can be induced by CRE recombinase were obtained from The Jackson Laboratory (Bar Harbor, ME). *Pstpip2*^*cmo*^ mouse strain was backcrossed on C57Bl/6J background for at least 10 generations and then used in the majority of experiments with the exception of experiments in Figure 1D and 4A, D, E, F, and S1. For these experiments the original *Pstpip2*^*cmo*^ strain on Balb/c genetic background has been selected due to the higher number of neutrophils that could be obtained by the negative selection method and due to the better quality of immortalized granulocyte progenitors derived from this strain. Both genetic backgrounds showed similar disease symptoms and similar dysregulation in ROS production. The BALB/c and C57Bl/6J inbred strains were obtained from the animal facility of Institute of Molecular Genetics, Academy of Sciences of the Czech Republic (Prague, Czech Republic). *Pstpip2*^*cmo*^-*DTA-MRP8-Cre* mouse strain was generated by breeding of the *Pstpip2*^*cmo*^ mice on C57Bl/6J background with *Gt(ROSA)26Sor*^*tm1(DTA)Lky*^/*J* strain mouse strain. Breeding this strain to *B6.Cg-Tg(S100A8-cre,-EGFP)1Ilw*/*J* mice carrying Cre transgene under the control of granulocyte-specific MRP8 promoter resulted in the genetarion of *Pstpip2*^*cmo*^*-DTA-MRP8-Cre* strain lacking almost all granulocytes (Supplementary Figure S4). Experiments in this work that were conducted on animals were approved by the Expert Committee on the Welfare of Experimental Animals of the Institute of Molecular Genetics and by the Academy of Sciences of the Czech Republic and were in agreement with local legal requirements and ethical guidelines.

### Primary Cells and Cell Lines

All primary cells and cell lines were cultured at 37°C with 5% CO_2_ in Iscove’s Modified Dulbecco’s Media (IMDM) supplemented with 10% fetal calf serum (FCS) and antibiotics. For bone marrow (BM) cell isolation, mice were sacrificed by cervical dislocation, BM was flushed with PBS supplemented with 2% FCS and erythrocytes were lysed in an ACK buffer (150 mM NH4Cl, 0.1 mM EDTA (disodium salt), 1 mM KHCO3). Murine neutrophils were isolated from BM cells using anti-biotin MicroBeads, and LS magnetic columns (Miltenyi Biotec, Bergisch Gladbach, Germany). For negative selection cells were labeled with biotinylated antibodies to B220, F4/80, DX5, c-Kit, CD3ɛ, CD115, and Ter119 prior to magnetic bead purification. For positive selection only anti-Ly6G-biotin was used. The purity of isolated cells was determined by flow cytometry. Primary murine monocytes were sorted from BM cells as Ly6G negative, Ly6C highly positive and Side-Scatter low cells using BD Influx sorter (BD Biosciences). The following cell lines were used in this study: HEK293FT cells (Invitrogen), Platimun Eco cells (Plat-E cells, Cell Biolabs, San Diego, CA), and immortalized granulocyte progenitors. For preparation of immortalized granulocyte progenitors we used a modified version of the protocol for generation of immortalized macrophage progenitors (Wang et al., 2006). The progenitors were first enriched by the depletion of Mac-1^+^, B220^+^, and Thy1.2^+^ from mouse BM cells and cultured in the presence of IL-3, IL-6, and SCF (supplied as culture supernatants from HEK293FT cells transfected with constructs coding for respective cytokines) for 2 days. Next, progenitors were transduced with ER-HoxB8 construct. The transduced cells were enriched for the GMP progenitor population by FACS (Lin^−^, Sca-1^−^, c-Kit^+^, FcγR^+^, CD34^+^) and propagated in a media containing 1 μM β-estradiol and 1% SCF-containing supernatant. Granulocyte differentiation was induced by β-estradiol withdrawal or by the β-estradiol withdrawal and replacement of SCF for G-CSF (50 ng/ml).

### Flow cytometry

Single cell suspensions of bone marrow cells were incubatecd with Fc-bloc and fluorophore-conjugated antibodies and analyzed on BD LSRII flow cytometer. For calcium response measurement, single cell suspensions of bone marrows from 6-8 week old mice were loaded with 2 μM calcium indicator Fura Red (Invitrogen). Samples were analyzed using a BD LSRII flow cytometer for 30 s at rest and then another 210 s after activation (either with fMLP, Silica or E.coli with OD=0.8). The relative calcium concentration was measured as a ratio of the Fura Red fluorescence intensity elicited by excitation wavelengths of 405 nm (emission measured at 635–720 nm) and 488 nm (emission measured at 655–695 nm). The data were analyzed with FlowJo software (TreeStar, Ashland, OR). Granulocytes were gated according to forward and side scatter properties.

### ROS meassurement

ROS production *in vitro* was assessed by luminol-based chemiluminescence assay as published previously (Goodridge et al., 2011). BM cells or purified murine neutrophils in IMDM supplemented with 0.2% FCS were plated in density of 10^6^ cells per well into a black 96-well plate (SPL Life Sciences, Naechon-Myeon, Korea). Cells were rested for 30 minutes at 37°C and 5% CO_2_. Then, luminol at final concentration 100 μM and stimuli (100 ng/ml LPS, fMLP 1 μg/ml, TNFα 10 ng/ml, E.coli OD_600_ ~ 0.8 – 5× diluted, Silica 50 μg/cm^2^, heat-agregated murine IgG 300 μg/ml, PMA 100 ng/ml) were added. Luminiscence was measured immediately on EnVison plate reader (Perkin Elmer, Waltham, MA); each well was scanned every minute for 70 minutes.

To assess ROS production *in vivo*, mice were intraperitoneally injected with luminescence reporter LC-012 in final concentration 75mg/kg (1,8mg/25g mouse) dissolved in PBS as previously described (Kielland et al., 2009). Luminiscence signal was acquired by Xtreme whole body imager (Bruker, Billerica, MA), with the following settings: binning 8×8, exposure time: 5 min. The quantification of photon counts was performed in Molecular Imaging Software (Bruker).

### DNA constructs

Generation of MSCV-PSTPIP2-EGFP construct. The coding sequence of mouse PSTPIP2 was amplified from cDNA of mouse myeloid progenitors (CMPs) and subcloned into pXJ41-EGFP cloning vector (Chum et al. 2016). IRES and Thy1.1 coding sequence was removed from MSCV-IRES-Thy1.1 retroviral vector (Clontech, Mountain View, CA) by digestion with EcoRI and ClaI followed by blunt ligation. PSTPIP2-EGFP coding sequence was then subcloned into modified MSCV vector using BglII and XhoI restriction sites to generate MSCV-PSTPIP2-EGFP.

Generation of MSCV-mPSTPIP2-TetOn inducible constructs. WT and mutated sequences (W232A or 3YF) of mouse PSTPIP2 described earlier (Drobek et al., 2015) were fused to C-terminal EGFP by PCR using P2A sequence as a linker. Fusion constructs were cloned into pLVX-Tet3G doxycycline inducible vector (Clontech) using AgeI and BamHI restriction sites. Resulting vectors were used as templates to amplify sequence spanning Tet-On 3G and TRE3G regulation/inducible elements together with PSTPIP2 by PCR and the resulting product was cloned it into MSCV-IRES-EGFP vector using ClaI and BglII restriction sites.

### Retroviral transduction

For confocal microscopy, c-kit+ stem and progenitor cells were obtained from BM of *Pstpip2*^*cmo*^ (C57Bl/6J) mice using magnetic purification (c-kit-biotin antibody, Anti-biotin mircobeads). Cells were expanded in IL-3, IL-6 and SCF supplemented media for 20h, then infected with PSTPIP2-EGFP retroviral construct. For the production of replication incompetent retrovirus, ecotropic packaging cells (Plat-E) were plated in 10 cm dish and transfected with 24 μg of plasmid DNA using Lipofectamine 2000 Reagent (Life Technologies) according to the manufacturer’s instructions. Virus containing supernatant was collected, concentrated with Amicon Ultra centrifugal filters with molecular weight cut-off 100 kDa (Merck Millipore) and immediately used to infect the cytokine expanded c-kit+ bone marrow cells. These cells were centrifuged with 150 μl of concentrated virus supernatant and 2.4 μl of Lipofectamine 2000 reagent (Sigma-Aldrich) at 1250 × g for 90 min at 30 °C and then incubated for another 4h at 37°C in 5% CO_2_ in a humidified incubator before the exchange of the media. Immortalized granulocyte progenitors were propagated in IMDM with 1 μM β-estradiol and 1% SCF-containing supernatant and then infected with PSTPIP2 mutant constructs using the same procedure described above.

### Microscopy

One day after infection, EGFP positive cells were sorted on Influx sorter and injected into sub-lethally irradiated (6 Gy in a single dose) CD45.1^+^ recipient mice. After 2 weeks, mice were sacrificed, BM cells isolated and resuspended in IMDM with 0.1% FCS. The cells were activated by 50 μg/cm^2^ fluorescent silica (see above) in a 96-well plate for 10 min. The cells were transferred to 4% paraformaldehyde in PBS and fixed at room temperature for 20 min. Cell nuclei were stained with 10 μg/ml Hoechst 33258 (Sigma) for 15 min. Cells were than washed 2x with PBS, resuspended in 150 μl of ddH2O and centrifuged on glass slide at 300 × g for 5min using Centurion Scientific K3 cytospin centrifuge (Centurion Scientific, Stoughton, U.K.). Cell samples were than mounted in 10 μl of DABCO mounting reagent (Sigma-Aldrich) and covered with glass coverslip (Zeiss, Oberkochen, Germany). Microscope setup: sequential 2-colour imaging was performed using a Leica TCS SP8 laser scanning confocal microscope (Leica, Wetzlar, Germany) with a 63×1.4 NA oil-immersion objective. Acquired images were manually thresholded to remove signal noise detected outside of the cell using ImageJ software.

### Cell activation, lysis, and immunoprecipitation

For Western blotting, the cells were washed and resuspended in IMDM with 0.1% FCS at a concentration of 1–4 × 10^7^ cells/ml. Subsequently, the cells were stimulated as indicated at 37°C. The activation of cells was stopped by the addition of an equal volume of a 2 × concentrated SDS-PAGE sample buffer (128 mM Tris pH 6.8, 10% glycerol, 4% SDS, 2% DTT) followed by the sonication and heating of the samples (99°C for 2 min). The samples were analyzed by SDS-PAGE followed by Western blotting. For detection of p47 phosphorylation, phosphoproteins were isolated from bone marrow cells using PhosphoProtein purification Kit (Qiagen, Hilden, Germany) according to manufacturer’s instructions, followed by detection of p47 by immunoblotting.

### RAC activity assay

2 × 10^7^ neutrophils (silica activated or not) were lysed in 1 mL Lysis buffer (25 mM HEPES pH 7.2, 150 mM NaCl, 10 mM MgCl_2_, 1 mM EDTA, 1% NP-40, 10% glycerol, 100 × diluted Protease Inhibitor Cocktail Set III (Calbiochem, Merck)) containing 5 μg PAK-RBD-GST (RAC-binding domain from PAK1 fused to GST, isolated from E.coli strain BL21 transformed with corresponding expression plasmid). After preclearing the lysate by centrifugation the complexes of active RAC and PAK-RBD-GST were isolated on Glutathion-Sepharose (GE Healthcare). RAC was then detected by immunoblotting.

### Anaesthesia

Mice for in-vivo imaging were anaesthetized by intramuscular injection of Zoletil (20mg/ml) – Xylazine (1mg/ml) solution with Zoletil dose 100mg/kg and Xylazine dose 1mg/kg.

### X-ray micro-computerized tomography

Hind paws were scanned in *in-vivo* X-ray μCT Skyscan 1176 (Bruker). Scanning parameters were: voltage: 50 kV, current: 250 μA, filter: 0.5 mm aluminium, voxel size: 8.67 μm, exposure time: 2 s, rotation step: 0.3° for 180° total, object to source distance: 119.271 mm, and camera to source distance: 171.987 mm with time of scanning: 30 min. Reconstruction of virtual slices was performed in NRecon software 1.6.10 (Bruker) with set-up for smoothing = 3, ring artifact correction = 4, and beam hardening correction = 36%. Intensities of interest for reconstruction were in the range from 0.0045 to 0.0900. For reorientation of virtual slices to the same orientation the DataViewer 1.5.2 software (Bruker) was used.

For μCT data analysis CT Analyser 1.16.4.1 (Bruker) was used. The volume of interest (VOI) was chosen there containing the distal part of hind paw starting from the half of metatarsus. Based on differences of X-ray absorption three parts were analyzed separately: the whole VOI, the newly formed bone connected mostly with arthritis, and the area inhabited by the original bone of phalanges and metatarsi. The total volume was recorded for all three parts. For original and new bone other parameters from 2D and 3D analysis were recoded to describe changes in the structure, namely: surface of the bone, surface-volume ratio, number of objects, closed porosity, mean fractal dimension, mean number of objects per slice, mean closed porosity per slice, and mean fractal dimension per slice. Scans with technical artifacts caused by spontaneous movements of animals were excluded from the analysis. Raw data are available upon request.

### Cytokine detection

Murine paws were homogenized in 1 mL RIPA lysis buffer (20 mM TRIS pH7.5, 150 mM NaCl, 1% NP-40, 1% Sodium deoxycholate, 0.1% SDS) containing 5 mM iodoacetamide (Sigma) and 100 × diluted Protease Inhibitor Cocktail Set III (Calbiochem, Merck) using Avans AHM1 Homogenizer (30s, speed 25). Any insoluble material was removed by centrifugation (20 000 × g, 5 min, 2°C) and concentration of the proteins in the samples were normalized to the same level using Bradford solution (AppliChem). Concentrations of IL-1β in the samples were determined by Ready-SET-Go!® ELISA kits from eBioscience according to the instructions of the manufacturer.

### Histology

The paws were fixed in 10% formol solution for 24h and decalcified in Osteosoft® (Merck) solution for 1 week, followed by paraffin embedding and histological cutting. The slides were stained in automatic system Ventana Symphony (Roche) and slides were scanned in Axio Scan.Z1 (Zeiss). The image postprocessing and analysis was done in Zen software (Zeiss).

### Statistical analysis

P values were calculated in GraphPad Prism software (GraphPad Software, La Jolla, CA) using unpaired t test (two-tailed) for data in Figure 2B-F, 5E; one-way ANOVA with Tukey-Kramer multiple comparison post test for data in Figure 3C, 4A, 6E, 7B, 7D; Mann-Whitney test for Figure 5C; and Gehan-Breslow-Wilcoxon test for disease-free curves in Figure 3D and 6C. Symbol meanings: n.s. P > 0.05; * P ≤ 0.05; ** P ≤ 0.01; *** P ≤ 0.001; **** P ≤ 0.0001. N numbers in figure legends represent number of biological replicates (in most cases animals).

## Acknowledgements

This work was supported by Czech Science Foundation (project 17-07155S), by institutional funding from the Institute of Molecular Genetics, Academy of Sciences of the Czech Republic (RVO 68378050), and by the Czech Centre for Phenogenomics (CCP, project no. LM2015040) and OP RDI CZ.1.05/2.1.00/19.0395 (Higher quality and capacity for transgenic models). We also acknowledge Light Microscopy Core Facility, IMG ASCR, Prague, Czech Republic, supported by MEYS (LM2015062), OPPK (CZ.2.16/3.1.00/21547) and NPU I (LO1419). JK and DG received additional support from Charles University Grant Agency (GAUK) (project number 923116).

