## Supplementary figures for "Regulation of neutrophil NADPH oxidase by PSTPIP2 affects bone damage in murine autoinflammatory osteomyelitis"

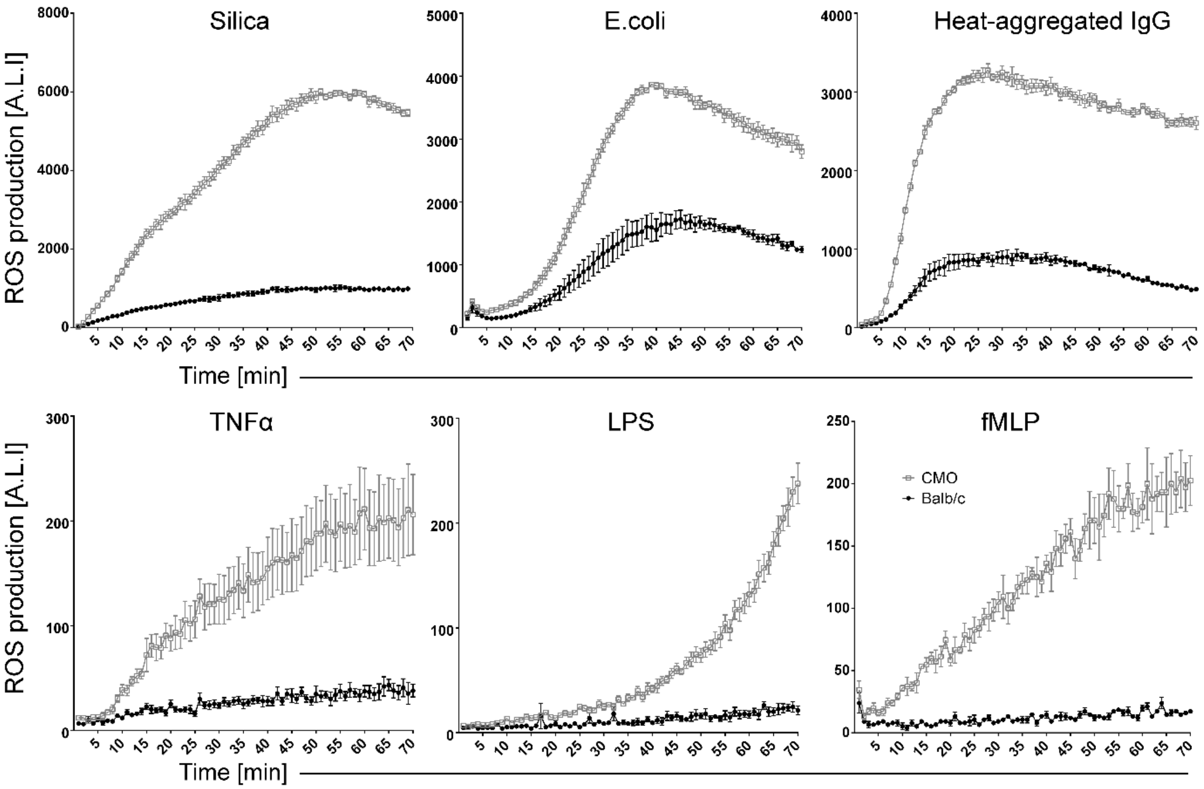


**Figure S1. Related to Figure 2. Dysregulated ROS production by bone marrow cells from *Pstpip2[cmo]* mice of Balb/c genetic background.** ROS production was detected by Luminol-based assay, exactly as described in Figure 2A.

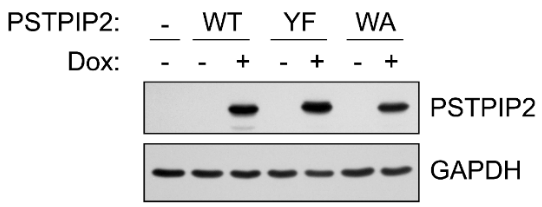


**Figure S2. Related to Figure 4. Verification of the expression of inducible PSTPIP2 constructs.**

*Pstpip2*$^{cmo}$ neutrophils differentiated from immortalized neutrophil progenitors retrovirally transduced with inducible expression constructs coding for PSTPIP2 wild-type (WT), PSTPIP2 W232A (WA), and PSTPIP2 Y323/329/333F (YF) were treated (or not) with Doxycycline (Dox) overnight and expression of PSTPIP2 was analyzed by immunoblotting. GAPDH was used as a loading control.

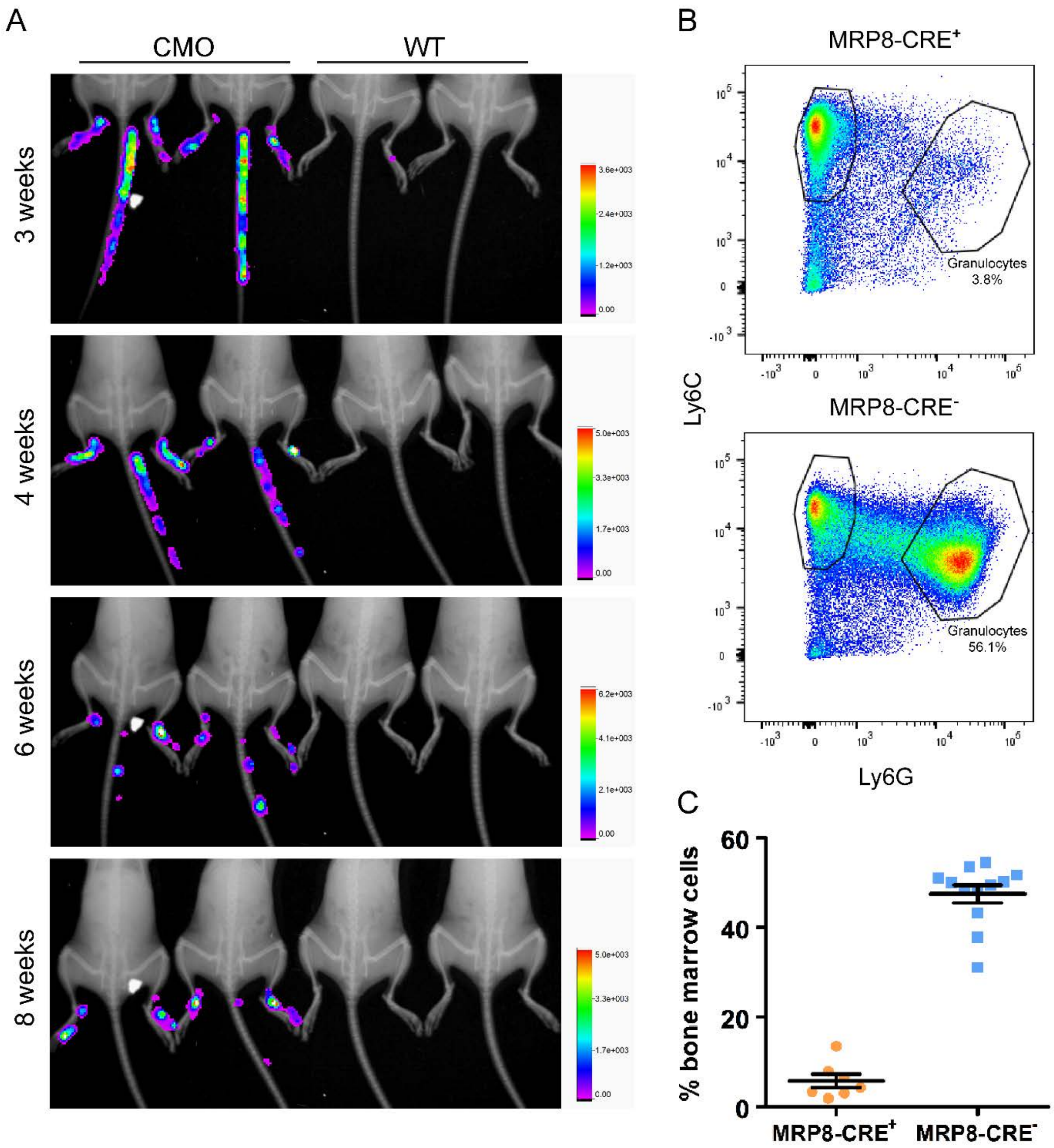


**Figure S3. Related to Figure 5. Neutrophil-dependent ROS production in vivo.**

(A) In vivo ROS production in WT and *Pstpip2^cmo^* (CMO) mice of various ages: representative images of the animals that were part of the analysis in Figure 5B.

(B) Neutrophils were depleted in Pstpip2$^{cmo}$-DTA mice via MRP8-CRE-dependent expression of Diphtheria toxin (MRP8-CRE$^{+}$) or left untouched in the absence of MRP8-CRE (MRP8-CRE$^{-}$). Representative FACS plots of CD11b$^{+}$ cells further gated for granulocytes (Ly6G$^{High}$, Ly6C$^{Low}$) and monocytes (Ly6G$^{-}$, Ly6C$^{High}$) are shown.

(C) Efficiencies of granulocyte depletion in multiple animals. Shown are the percentages of granulocytes gated as in (B).

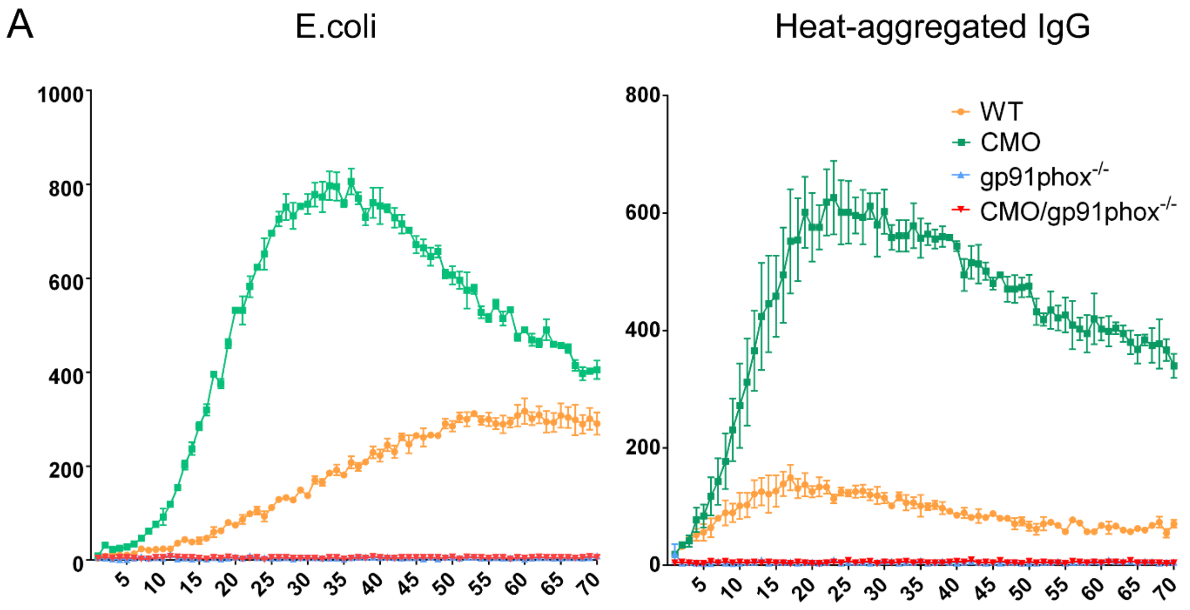


**Figure S4. Related to Figure 6. Absence of ROS production in bone marrow cells from *gp91phox$^{-/-}$* and *Pstpip2$^{cmo}$/gp91phox$^{-/-}$* mice.**

BM cells from mice of indicated genotypes were activated by live E.coli bacteria or heat-aggregated IgG in the presence of luminol and ROS-dependent luminescence was quantified in 1 min intervals.

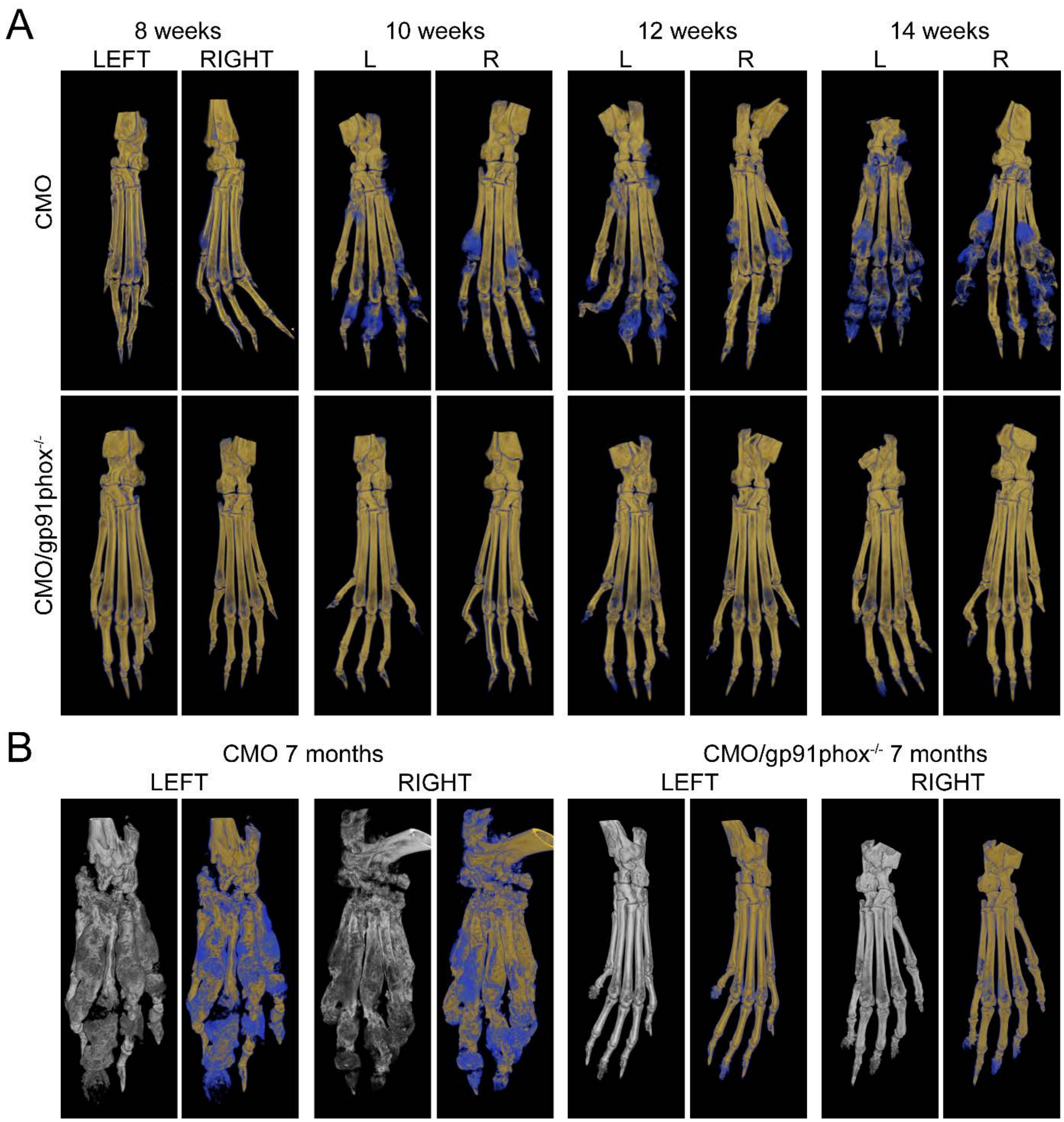


**Figure S5. Related to Figure 7. Time course of bone damage development in *Pstpip2$^{cmo}$* and *Pstpip2$^{cmo}$/gp91phox$^{-/-}$* mice.**

(A) Mice at 8, 10, 12, and 14 weeks of age were anesthetized and their hind paw bones were imaged on X-ray µCT scanner.

(B) The same analysis performed on 7 months old animals.

Grey images represent visualization of total bone tissue. Pseudocolor images distinguish between old (in yellow) and newly formed (in blue) bone mass.
